# EVOLVE: A Web Platform for Evolutionary Phase Analysis and New Variant Exploration from Multi-Sequence Data

**DOI:** 10.1101/2024.12.26.630381

**Authors:** Satyam Sangeet, Anushree Sinha, Madhav B. Nair, Arpita Mahata, Raju Sarkar, Susmita Roy

## Abstract

Recurrent updates from the World Health Organization (WHO) on refining definitions for Variants of Concern (VOCs), Variants of Interest (VOIs), and Variants Under Monitoring (VUMs) underscore the need for systematic investigations to develop quantifiable metrics that differentiate critical phases of variant evolution. This study illustrates how protein data can be systematically analyzed by applying phase transition principles from statistical mechanics, where quantifying mutational response function (MRF) helps identify concerning variants along evolutionary paths. To support the exploration of metrics like mutational entropy, MRF, and other relevant indicators for novel or uncharacterized viruses and pathogenic bacteria, we introduce EVOLVE—a web platform that empowers researchers across diverse fields to access sequence-based analyses. EVOLVE streamlines data upload and analysis with a user-friendly interface and comprehensive tutorials. Overcoming the challenges with large sequence-space analysis its AI-driven data-learning module integrates evolutionary dynamics with microscopic mutational information, enabling the prediction of future site-specific mutations in prospective missense variants. Access EVOLVE for free at https://www.evolve-0fm3.onrender.com.

## Introduction

The intricate relationship between an organism’s ecological and evolutionary dynamics is shaped by its rapid mutation. RNA viruses, in particular, have very high rates of mutation because they do not have a mechanism for proofreading during replication^1^. As a result, it is easier for various viral variants to arise, which helps them avoid host immune reactions and adapt to the changes in environment. Recombination is another process that certain RNA viruses go through to broaden their genetic diversity and produce new viral lineages that may have different phenotypes^2^. The recent COVID-19 pandemic vividly demonstrated this evolutionary pattern, showcasing how the SARS-CoV-2 RNA viruses rapidly acquire mutations and recombine, potentially leading to more infectious and transmissible variants^3,4,5^.

Because of this ongoing evolution, in late 2020, the emergence of the SARS-CoV-2 variants that increased global public health risks led the World Health Organization (WHO) to classify some as variants of interest (VOIs) and variants of concern (VOCs)^6^. This was done to prioritize global monitoring and research and to inform and adjust the COVID-19 response. The US Centers for Disease Control and Prevention (CDC) and the World Health Organization (WHO) categorize these variants into three main groups: Variants Being Monitored, Variants of Interest, and Variants of Concern^6^. Variants are classified as “of interest” when they are linked to community transmission and genetic changes. They are deemed “of concern” if they exhibit increased transmissibility, virulence, or reduced effectiveness of health interventions. Along with monitoring community transmission of SARS-CoV-2, it also remains essential to closely monitor genetic changes effectively, which are crucial aspects of the global strategy to address the occurrence of mutations that have negative public health implications. Even subtle alterations in genetic sequence can have far-reaching effects, influencing transmissibility, virulence, diagnostic accuracy, vaccine efficacy, antiviral treatment strategies, and the effectiveness of public health and social interventions^7,8,9,10,11,12^. To address this, WHO recently recommended close monitoring of genetic changes in SARS-CoV-2 variants that are suspected to influence viral characteristics or exhibit early signs of a growth advantage over other circulating variants. In such instances, where the phenotypic or epidemiological impacts of these changes are uncertain, such variants are categorized as ‘Variants Under Monitoring’ (VUM). In October 2023, WHO further updated the VUM definition where they include variants with many mutations in antigenic sites, along with community transmission detected in more than two countries within two to four weeks^13^. These frequent updates on their definition clearly reflect that this classification is somewhat qualitative, lacking precise and consistent metrics.

On another account, viral evolutionary dynamics has experienced a paradigm change. Moving from theoretical and qualitative studies^14,15^ to large-scale genomic analyses of human pathogens^16,17,18^ this development is fueled by three major advancements in the field: (I) Establishment of big data infrastructures, such as NCBI Virus (https://www.ncbi.nlm.nih.gov/labs/virus/vssi/<u>#/</u>) and GISAID (https://www.gisaid.org/^19,20,21^), which allows the researchers to easily access and analyze the huge genomic resources (II) Advancements in computational hardware allows large-scale genomic analysis due to the efficient execution of complex algorithms and (III) Advanced statistical techniques have been developed to analyze the evolution and transmission of viruses, such as phylogenetic analysis^22^, network analysis^23^, information theory^24^, and population genetics^25^. While databases like NCBI and GISAID indeed provide extensive viral sequence repositories, they primarily offer raw data with limited annotations on evolutionary trajectories. Beyond basic phylogenetic analysis, these resources provide minimal contextual information regarding how specific mutations influence viral fitness, transmissibility, or adaptability over time. Again, while various computational algorithms for network analysis^23^, population genetics^25^ have improved our ability to study evolutionary dynamics at a large scale, predicting mutational changes of a variant and their molecular impacts —particularly those that have potential to alter protein structure or function— remains a significant challenge. Although computational tools can model structure-based molecular effects of point mutations using ΔΔG calculations^26^, they operate independently, creating a disconnect between structural predictions and broader evolutionary processes. For example, a typical viral protein with 300 residues can undergo 5,700 possible single amino acid substitutions, and this mutational landscape becomes exponentially more complex when multiple mutations are considered. The sheer scale of possibilities renders comprehensive computational analysis infeasible, creating a bottleneck in our ability to predict and understand viral evolution. Without a systematic framework to prioritize significant mutations, researchers face the challenge of navigating an unmanageable mutational phase space or relying on educated guesses, which risk missing critical evolutionary insights.

Despite these challenges, studies of viral evolution have made important contributions, such as tracing the origins and spread of HIV^27,28,29^ and uncovering the roles of natural selection and migration in shaping influenza A dynamics^30,31,32^. However, ongoing research on SARS-CoV-2 variants highlights the need to understand viral evolution systematically, connecting microscopic information regarding mutation prone sites on a protein structure and evolutionary dynamics over large sequence space^33^. In this context, statistical mechanics provides a framework to capture the collective response of mutations over large genomic sequence space, while machine learning serves as a processing tool for such vast genomic data analysis, which can further be used for attaining microscopic information such as site-specific mutation prediction or collection of mutations that may again give rise to a drastic change in the evolutionary dynamics. These approaches combined can be potentially utilized to enhance the understanding of viral adaptation and spread which is crucial for mitigating risks from emerging strains.

Building on this need, we present EVOLVE (**<u>E</u>**volutionary **<u>V</u>**ariant **<u>O</u>**bservation, **<u>L</u>**earning and new-**<u>V</u>**ariant **<u>E</u>**xploration), a web platform integrating statistical mechanics and machine learning to analyze viral evolution. Built on extensive analysis of the SARS-CoV-2 surface glycoprotein data, EVOLVE integrates protein data analysis, offering advanced evolutionary insights. Successfully tested on several species such as SARS-CoV-2, Influenza, and *Xanthomonas*, EVOLVE demonstrates versatility across diverse species. With a user-friendly interface, accompanied by a detailed, step-by-step tutorial for guidance, EVOLVE democratizes viral evolution studies, providing a deeper understanding and improved management of viral threats. EVOLVE is freely accessible at www.evolve-0fm3.onrender.com.

## Results and Discussion

### Interface

EVOLVE is equipped with an intuitive graphical user interface compatible with all modern web browsers, ensuring seamless accessibility for researchers. Upon registration, users are granted access to a personalized account that centralizes all necessary computational tools and resources. The platform facilitates effortless data upload through dedicated calculation pages, allowing users to submit their input files efficiently. Once the data is processed, EVOLVE generates comprehensive results, which are made available for download via convenient links. The results are meticulously presented in a clear and interpretable format, enabling users to easily grasp the visualized outcomes. This streamlined workflow exemplifies EVOLVE’s commitment to providing a robust and accessible environment for advanced data analysis on a web-based platform.

### EVOLVE Workflow

The EVOLVE pipeline (Fig. 1) describes an organized method for analyzing data for both genomic and amino acidic sequence space. The workflow first begins with preprocessing the data extracted from different resources followed by several critical property calculations discussed as follows:

**Fig. 1:**
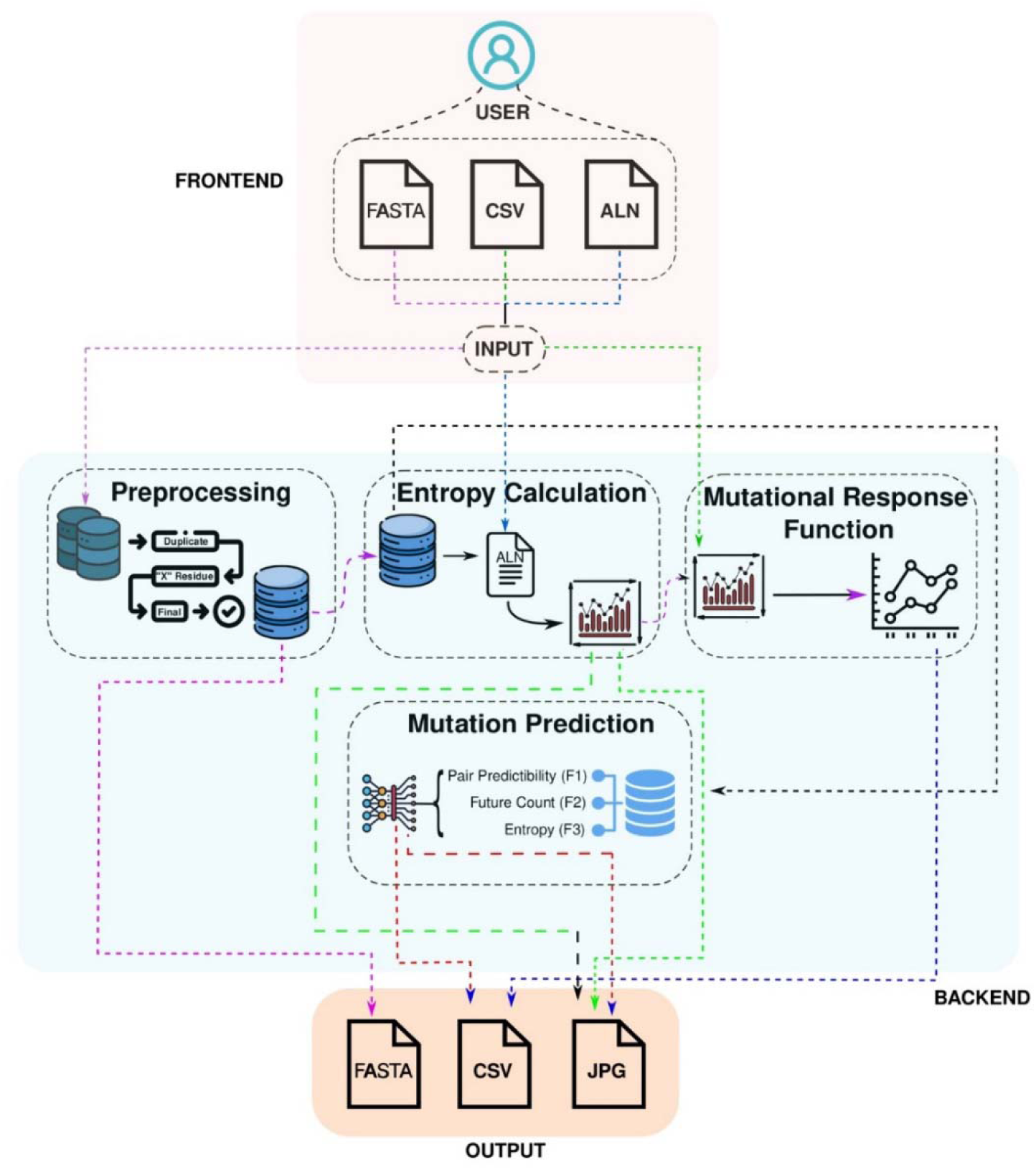
Overview of EVOLVE web server. The front end allows the user to upload files in the specified format. The backend performs the necessary calculations selected by the user. CSV: comma-separated values; ALN: MSA alignment file

#### Preprocessing

In the workflow, Preprocessing is the first step in the process of refining the dataset. It involves several crucial steps. The protein *fasta* files first go through preprocessing that results in the removal of the entries that contain “**X**” which correspond to undefined amino acids within the sequences, as well as removing duplicate sequences. The thorough preprocessing guarantees that unwanted or incorrect entries that could jeopardize the integrity of the dataset are excluded. The preprocessed data is then used for performing Multiple Sequence Alignment (MSA) and the results are obtained in *.aln* (Alignment) format. The MSA procedure aligns the sequences and prepares them for further analyses in EVOLVE. It can be carried out with other preferred MSA software such as MAFFT^34^ and ClustalW^35^.

### Mutational Entropy Calculation

Following the preprocessing stage, the next critical step in the EVOLVE workflow is the calculation of mutational entropy, specifically utilizing Shannon entropy as the foundational metric. Shannon entropy quantifies the uncertainty or variability of amino acid residues at each position within the aligned sequences, providing a robust framework for understanding the evolutionary dynamics at play. This calculation involves assessing the frequency distribution of each amino acid at a given position, allowing for the determination of the degree of variability present. High Shannon entropy values indicate a diverse array of residues, suggesting that the position may be subject to relaxed selective pressure, thereby allowing for greater mutational freedom. Conversely, low entropy values reflect a predominance of specific residues, indicating strong evolutionary constraints that are likely essential for maintaining the structural and functional integrity of the protein^36,37^. Evaluating the mutational entropy calculation, researchers can effectively identify mutational hotspots—regions of the protein that are more prone to variation and may play critical roles in adaptive evolution.

### Analysis of Mutational Response Function to Explore Evolutionary Phases

The analysis of the Mutational Response Function (MRF) represents an important advancement in understanding the evolutionary trajectories of viral variants. The MRF is designed to capture mutational state transitions by quantifying the mean value of mutational entropy alongside its mean square fluctuations. By analyzing these metrics, critical transitions can be identified in the evolutionary process, providing insights into how viral populations adapt over time. The mean value of mutational entropy reflects the overall variability within the sequence data, while the mean square fluctuation offers a measure of the stability of this variability. Together, these parameters discern patterns of mutational behavior that may indicate significant evolutionary shifts. This approach not only enhances the understanding of the dynamics of viral evolution but also facilitates the identification of key moments in the evolutionary timeline where substantial changes may occur.

#### New Mutation Prediction Employing Statistical Mechanics Integrated AI-based Method

The EVOLVE platform incorporates a Multi-Layer Perceptron (MLP) model that is trained on extensive datasets, enabling it to predict potential mutational residues with high accuracy. By analyzing features such as pair predictability of amino acids, future composition, and entropy calculations, the MLP model generates predictions that highlight residues with the highest likelihood of undergoing significant mutations. This predictive capability is particularly valuable in the context of emerging viral variants, as it allows us to focus on the potential residues that have a significant probability of mutating in the future.

#### Brief Tutorial with Illustrative Examples

A detailed tutorial about the usage of the EVOLVE server is provided on the website: https://evolve-0fm3.onrender.com/tutorial. However, through illustrative examples, we demonstrate the usage of EVOLVE by analyzing a total of four cases: three cases of viral species: Omicron variant of SARS-CoV-2, Hemagglutinin protein of H1N1 Influenza virus and Coat Protein of Citrus Tristeza Virus, and a Bacterial species *Xanthomonas oryzae*. Additionally, we provide the Transfer Learning method, which enables the user to use the previously trained model and retrain it using fresh data, utilizing the Dengue virus’s NS3 protein and Ebola virus’s glycoprotein as an example.

### Case I: Omicron Variant

#### Data Collection and Preprocessing of Evolutionary Data

Taking advantage of the protein data, the evolutionary history and mutational localization of the Omicron variant were examined using EVOLVE. The raw FASTA files comprising protein sequences of the surface glycoprotein of the Omicron variant (B.1.1.529), were obtained from the NCBI Virus database (https://www.ncbi.nlm.nih.gov/labs/virus/vssi/#/) in a month-wise manner. Multiple Sequence Alignment (MSA) was performed using a refined dataset of month-wise sequences obtained through subsequent preprocessing and thereby, a month-wise evolutionary trajectory was built. An analysis of the evolutionary trajectory from September 2021 to February 2022 was done for cumulative protein entropy.

#### Mutational Entropy Calculation

To calculate site-specific entropy, alignment files were processed using the EVOLVE server, revealing notable evolutionary trends. In B.1.1, the N-terminal domain (NTD) exhibited fluctuating entropy between August and November 2021, followed by a sharp decline in December 2021 (Fig. 2a, green). This reduction suggests the fixation of specific mutations within the viral population, marking a critical evolutionary event^38^. The entropy decline indicates a stabilization phase in the NTD, where advantageous mutations, likely driven by immune escape or selective pressures, became fixed in genome^39^. This behavior aligns with the evolutionary processes in other species, where reduced genetic variability reflects the emergence of fitter variants^40,41^.

**Fig. 2:**
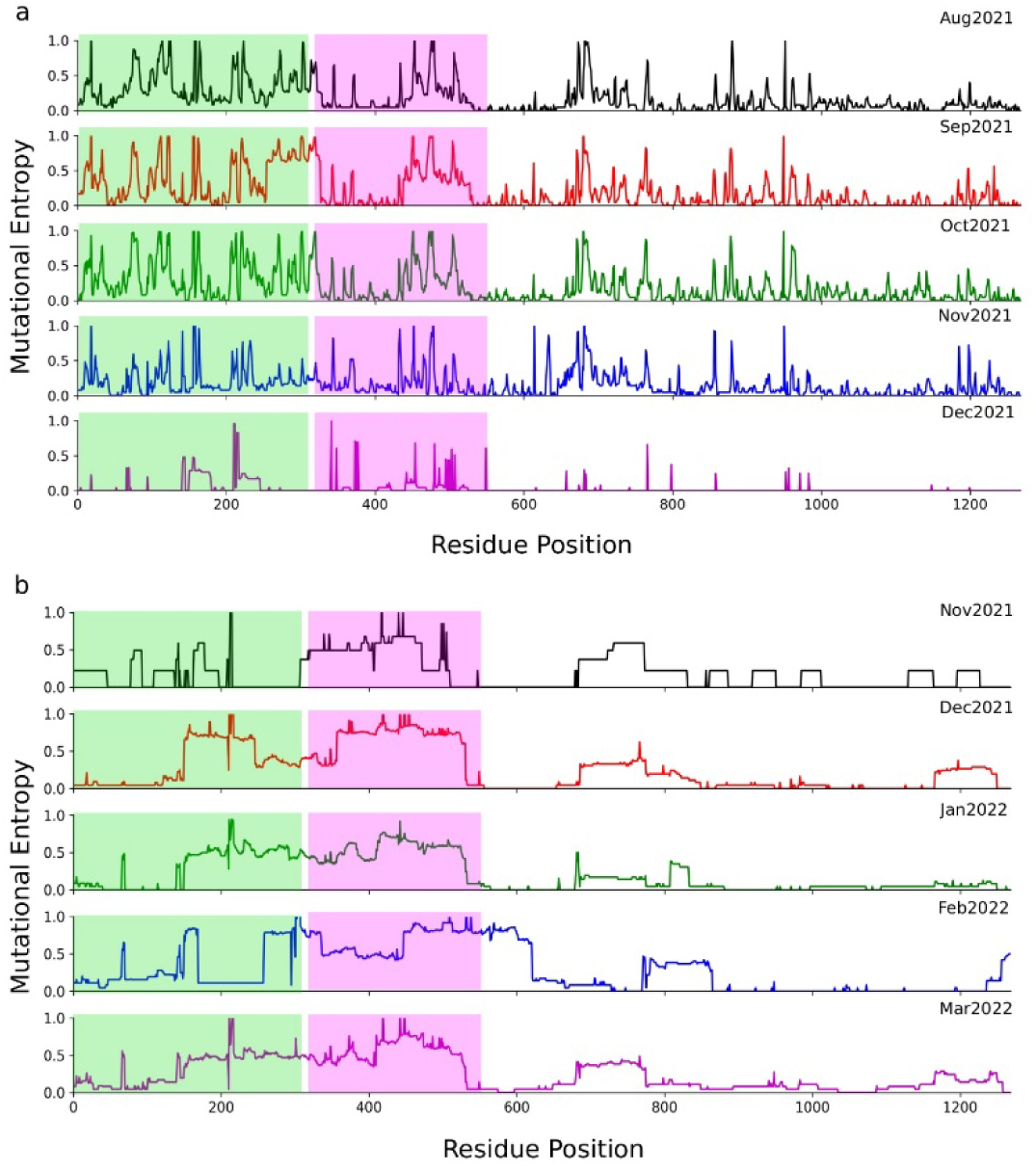
Mutational Entropy of (a) B.1.1. and (b) Omicron variant of SARS-CoV-2. The graph represents the time-dependent entropic variation in the protein sequence of the viruses. The green-shaded region corresponds to the N-terminal domain (NTD) and the magenta-shaded region corresponds to the receptor binding domain (RBD) of the spike glycoprotein.

Conversely, in Omicron, from December 2021 through March 2022, NTD entropy increased significantly, indicating a phase of diversification as the virus explored a broader mutational space, likely in response to immune pressure (Fig. 2b, green). This elevated entropy suggests ongoing selection for immune-evasive mutations, highlighting a dynamic period in Omicron’s evolution. Such rapid diversification mirrors adaptive responses to environmental changes^42,43^. In the receptor-binding domain (RBD), entropy patterns further distinguish B.1.1 from Omicron. In B.1.1, RBD entropy remained high through November 2021, indicating ongoing exploration of mutations to optimize ACE2 receptor binding and immune evasion (Fig. 2a, magenta). In Omicron, a pronounced increase in RBD entropy from December 2021 suggests the emergence of advantageous mutations, sparking diversification (Fig. 2b, magenta). These shifts illustrate Omicron’s adaptive evolution, like bursts of phenotypic variability seen in species under strong selective pressure^44^. These entropy patterns highlight key evolutionary events in SARS-CoV-2 adaptation, correlating B.1.1’s entropy decline with Omicron’s emergence and subsequent mutational exploration.

#### Evolutionary Phase Analysis and Observation of Dynamic Phase Transition in the case of Omicron

EVOLVE facilitates access to the evolutionary and dynamic phase transitions along evolutionary trajectories by calculating the Mutational Response Function (MRF)^45^. Monthly entropy data was uploaded to the EVOLVE server for MRF calculation. The analysis revealed distinct evolutionary patterns between the Omicron and B.1.1 variants. The mutational entropy of the Omicron surface glycoprotein (Fig. 3a, blue) was significantly elevated compared to B.1.1 (Fig. 3a, red) during November-December 2021 timeframe, indicative of heightened mutational variability and exploration. In contrast, the mutational entropy of Omicron NTD (Fig. 3b, blue) was lower than that of B.1.1 (Fig. 3b, red), suggesting a phase of increased stabilization in this critical domain. Interestingly, the B.1.1 variant consistently exhibited lower mutational entropy compared to Omicron in both the surface glycoprotein and NTD regions during the emergence of the Omicron variant. Moreover, the MRF for the Omicron surface glycoprotein (Fig. 3c, blue) showed a pronounced spike, exceeding that of B.1.1 (Fig. 3c, red), in the November-December 2021 window. This pattern implies an initial period of mutational stabilization of the B.1.1 lineage, followed by a surge in evolutionary exploration coinciding with the emergence of Omicron^46,47^, underscoring the MRF’s utility in signaling significant evolutionary events. Notably, the MRF for the B.1.1 variant was also higher during this critical transition period, corresponding with the elevated MRF observed in the Omicron surface glycoprotein. Furthermore, the Omicron NTD (Fig. 3d, blue) also exhibited a slightly elevated MRF compared to B.1.1 (Fig. 3d, red) during the November-December 2021 timeframe, albeit to a lesser degree than the surface glycoprotein. These findings suggest that the MRF may be a critical parameter in understanding the phase transitions experienced by SARS-CoV-2 during the emergence of new variants.

**Fig. 3:**
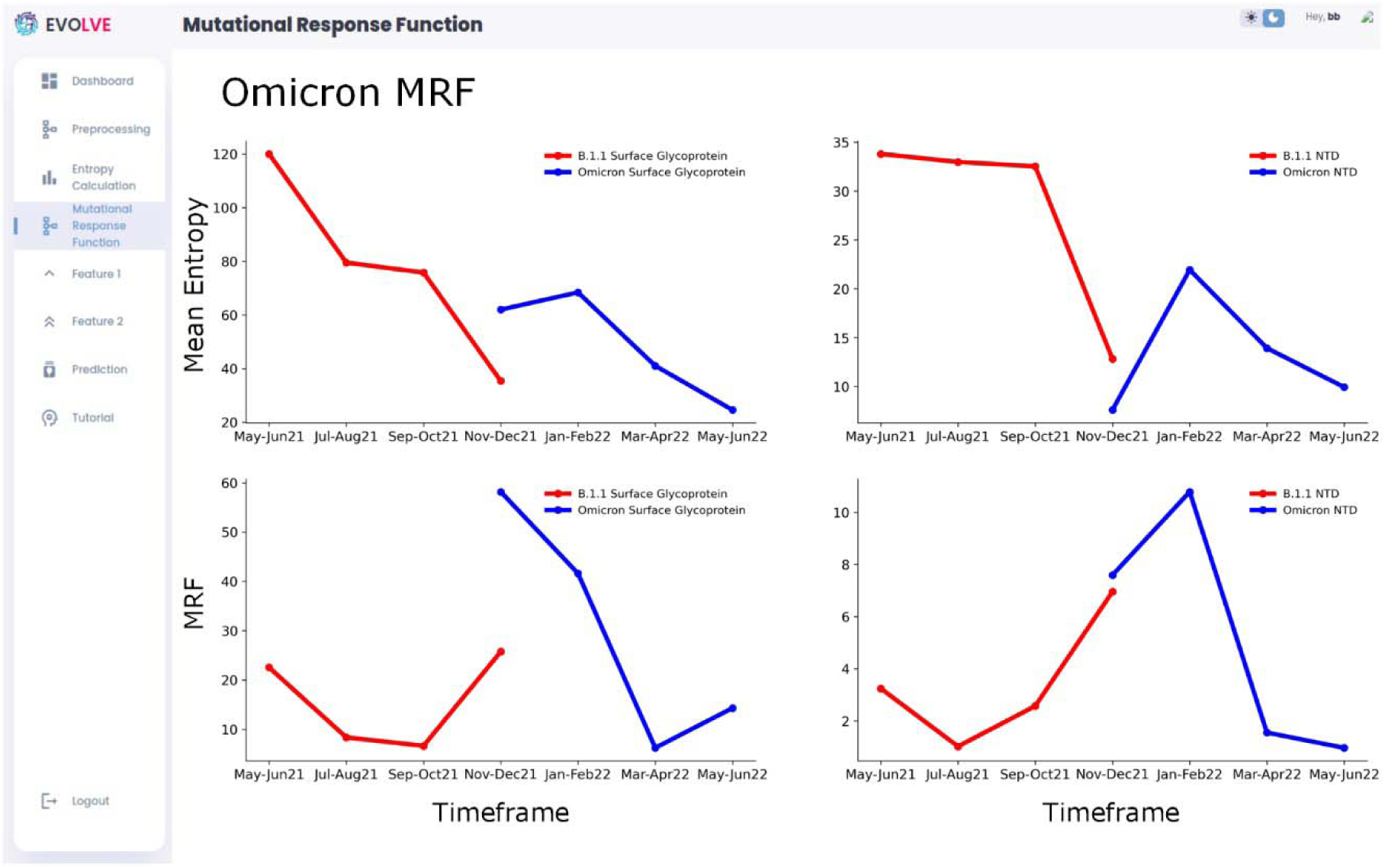
Mean Entropy and Mutational Response Function for Surface Glycoprotein and N-Terminal Domain of the B.1.1 and Omicron variants. (a) Mean Entropy of B.1.1 (red) and Omicron variant (blue) (b) Mean entropy of NTD of B.1.1 (red) and Omicron (blue) variant (c) MRF of B.1.1 (red) and Omicron (blue) variant (d) MRF of NTD of B.1.1 (red) and Omicron (blue) variants.

This analysis underscores a key principle: monitoring MRF and entropy can predict the emergence of descendant variants. The sudden increase in B.1.1’s MRF preceded Omicron’s appearance, while Omicron’s initial high MRF and subsequent decline exemplify a new variant’s typical pattern of initial diversity followed by stabilization^48^. Continuous tracking of MRF and entropy in dominant strains may provide early warning of significant evolutionary shifts, enhancing pandemic surveillance and response strategies.

Building upon our insights from the MRF and entropy analyses, we developed a machine learning approach to predict specific mutational sites in the Omicron variant. After optimizing our model through feature selection (Supplementary Fig. S1), we implemented a Feedforward Backpropagation Multilayer Perceptron (MLP) with three input features, two hidden layers of eight neurons each, and a binary output predicting mutation probability (Fig. 4a). Crucially, the model was trained exclusively on sequences from the original Omicron variant, withholding all data on Omicron sub-variants. When predicted on these sub-variants, the model identified 13 potential mutation sites (Fig. 4b). Four predicted sites (Table 1): Asparagine at position 334 (N334), Serine at position 494 (S494), Alanine at position 653 (A653), and Threonine at position 791 (T791) had already acquired mutations in circulating Omicron sub-variants (Fig. 4c, d Supplementary Table S1). These spike residues play critical roles in SARS-CoV-2’s infectivity, immune evasion, and adaptability, underscoring the accuracy of model predictions.

**Fig. 4:**
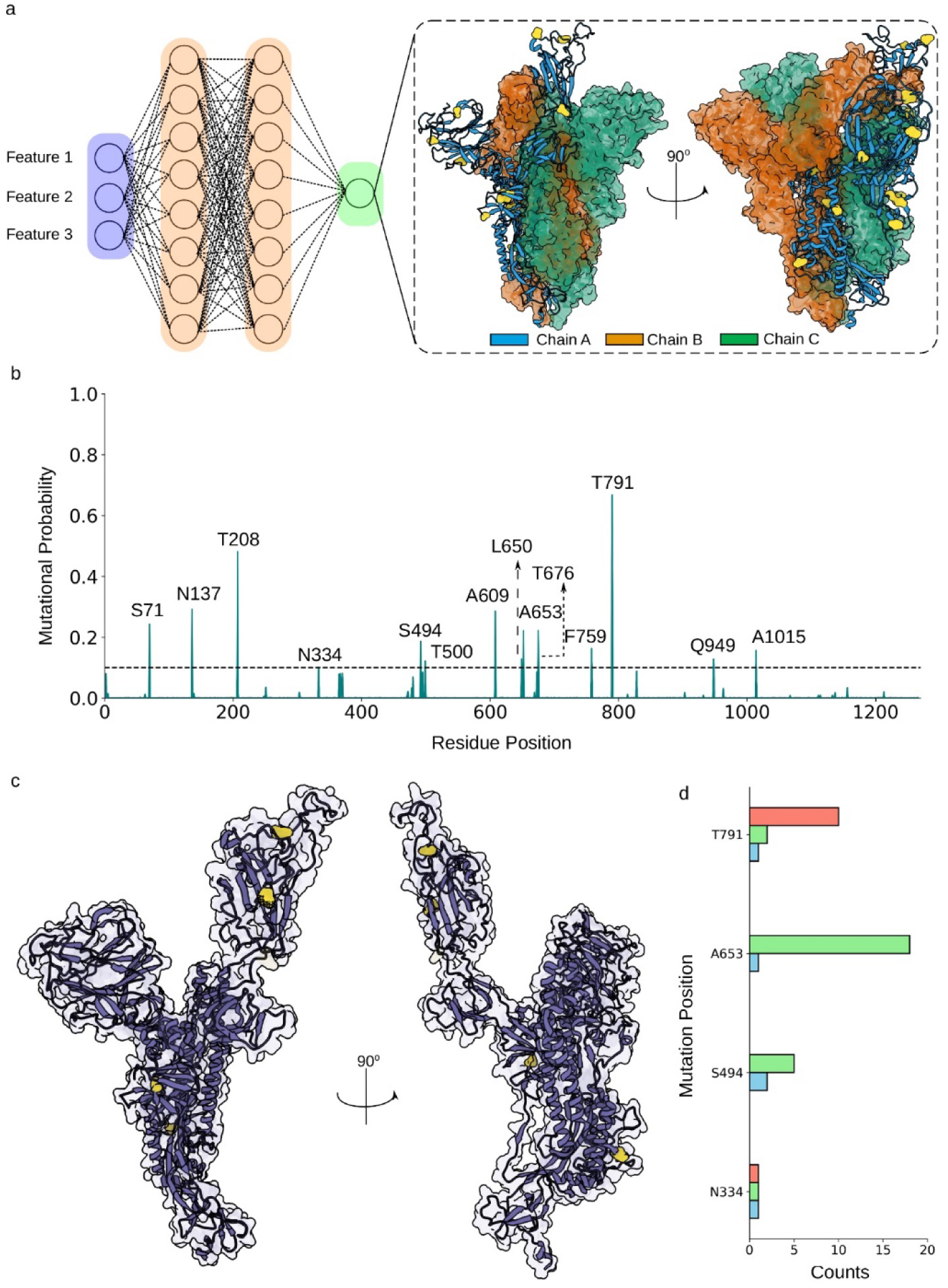
EVOLVE analyses for Omicron variant. (a) MLP architecture for the mutation prediction in the spike protein of the Omicron variant. The inline image shows the surface view of the trimeric spike protein with predicted mutating residue (yellow) according to the model prediction. (b) Mutational probability graph showing the model prediction. The cutoff of 0.1 is a user-defined cutoff suggesting that residues showing a mutational probability of more than 0.1 have a higher chance of mutating in the near future and vice versa. (c) Single chain aptamer of spike protein with the four major predicted mutating residues that mutate in the sub-variants of Omicron. (d) Bar plot showing the frequency of major predicted residues mutating into other amino acids in different variants. The details about the variants are provided in the supplementary file.

**Table 1:** Model prediction for mutating residues in Omicron and Influenza virus which are shown to be relevant for the viral survival and are experimentally validated.

| Omicron Mutation Prediction |  | Influenza Mutation Prediction |  |
| --- | --- | --- | --- |
| Position | Residue | Position | Residue |
| 334 | Asparagine | 145 | Lysine |
| 494 | Serine | 158 | Glycine |
| 653 | Alanine | 186 | Serine |
| 791 | Threonine | 223 | Valine |

##### N334

Asparagine at position 334 makes an important hydrogen bond with S309 antibody. This interaction of antibody, along with N334 of RBD, supports the antibody’s capacity to neutralize the Omicron variant, and plays a major role in structural stability and binding affinity^49^. This epitope’s stability, which is primarily preserved by N334 and nearby residues, guarantees that S309 can neutralize the virus efficiently even when Omicron develops mutations. N334 is an N-glycosylation site, and mutations that inhibit glycosylation at this site are known to decrease viral infectivity, further demonstrating the functional significance of this position in immune response modulation and viral transmission.

##### S494

The S494P mutation enhances immune escape without compromising the spike protein’s ability to bind ACE2. By conferring partial resistance to neutralization by convalescent sera, S494P has been observed in several highly transmissible variants^50^. The positive selection of this mutation suggests a fitness advantage, particularly when combined with immune-evasive mutations like E484K and N501Y, which together amplify immune resistance. S494P’s role in evading neutralizing antibodies while supporting effective ACE2 binding underlines its importance in viral transmission and immune evasion.

##### T791

T791, which is in the heptad repeat region of the spike protein, is significant because the T791I mutation affects viral replication and immune evasion by disrupting a phosphorylation site. This mutation, which enables the virus to adapt and endure in a host environment, is commonly seen in chronic infections, especially in immunocompromised patients^51^. T791 is a crucial location for comprehending viral adaptability since it can change the activity of spike proteins and possibly change the virus’s behavior under immunological pressure by modifying phosphorylation.

The precision and resilience of the model are demonstrated by its capacity to forecast these mutation-prone residues, all of which are essential for SARS-CoV-2’s infectivity, immunological evasion, and adaptability. By pinpointing these crucial locations, the model demonstrates its usefulness as a prediction tool for monitoring and comprehending new variations. This alignment between the model’s predictions and observed mutations in variants demonstrates its capacity to identify key sites of potential mutation. To assess the functional implications of these mutations, we conducted stability analysis using the mCSM^26^ webserver. Most potential mutations at the four key sites were predicted to be destabilizing, with a subset conferring increased stability (Supplementary Fig. S2). Notably, stabilizing mutations have not yet been observed in circulating variants, suggesting a potential evolutionary path for future variants.

Entropy analysis reveals numerous mutational hotspots in SARS-CoV-2 spike protein (Fig. S3a), but the sheer volume of high-entropy residues complicates prioritization for further study. Our ML approach addresses this challenge by efficiently screening these residues to identify those with the highest potential impact on protein structure and function (Fig. S3b). Trained on extensive datasets, the ML model discerns patterns and correlations often overlooked by conventional methods, effectively distinguishing signal from noise in entropic data. While entropy analysis identifies multiple mutation-prone regions, our ML approach refines this to crucial single-point mutations, which often have more significant functional or stability impacts. While SARS-CoV-2 benefits from an extensive genomic dataset, many pathogens lack such comprehensive sequencing data, making evolutionary trajectory predictions challenging. Nevertheless, the approach remains valuable for identifying potential mutational sites across diverse species. We demonstrate this versatility by applying EVOLVE to three distinct cases: Influenza virus, *Xanthomonas oryzae*, and Citrus tristeza virus (CTV). In each instance, despite limited data sets compared to SARS-CoV-2, EVOLVE successfully identified key mutational sites, some of which align with known variants or functionally important regions. This cross-species applicability underscores EVOLVE’s potential as a universal tool for predicting mutational hotspots in various pathogens.

### Case II: H1N1 Influenza virus

As a second case study, we analyzed the Hemagglutinin (HA) protein of the H1N1 influenza virus (Fig. 5a), which encompasses critical functional elements including receptor-binding domain and antigenic epitopes (Fig. 5b). Previous research indicates that, in the context of influenza, positive selection operates inefficiently at the level of individual hosts, with stochastic processes predominantly shaping host-level viral evolution^52^. This suggests that the intra-host evolution of the influenza virus is relatively slow and largely governed by genetic drift and other random processes, rather than by positive selection for antigenic variants^52^. Additionally, the reassortment rate of influenza is constrained during human infection, implying that the virus may evolve less rapidly than previously assumed^53^. EVOLVE predicted 26 potential mutation sites in the Hemagglutinin (HA) protein of H1N1 influenza virus (Fig. 5c, d). Several of these predicted positions align with experimentally validated mutations of significant functional importance (Table 1). Notably, predicted positions 145 and 158 correspond to experimentally observed mutations K145E^54^ and G158E^54^, which are associated with antibody escape in the Ca2 and Sa antigenic sites, respectively. The model successfully identified position 186, which has been implicated in both B-cell antigenic recognition and host species transition from avian to human^55^. Furthermore, position 223, another predicted site, has been experimentally confirmed as crucial for receptor-binding properties^55^. Several predicted mutations (positions 145, 186, and 223) are in functionally critical regions: antigenic sites, receptor-binding domains, and host adaptation sites^56^. These findings demonstrate EVOLVE’s capability to identify evolutionary significant positions within viral proteins. The model’s predictions encompass residues involved in immune escape (antigenic sites), host adaptation, and receptor binding, suggesting its utility in anticipating functionally relevant mutations. Furthermore, the analysis revealed that, except for seven mutations, most of the predicted mutations are associated with destabilizing effects on the protein (see Supplementary Fig. S4). Detailed list of mutations is provided in Supplementary table S2.

**Fig. 5:**
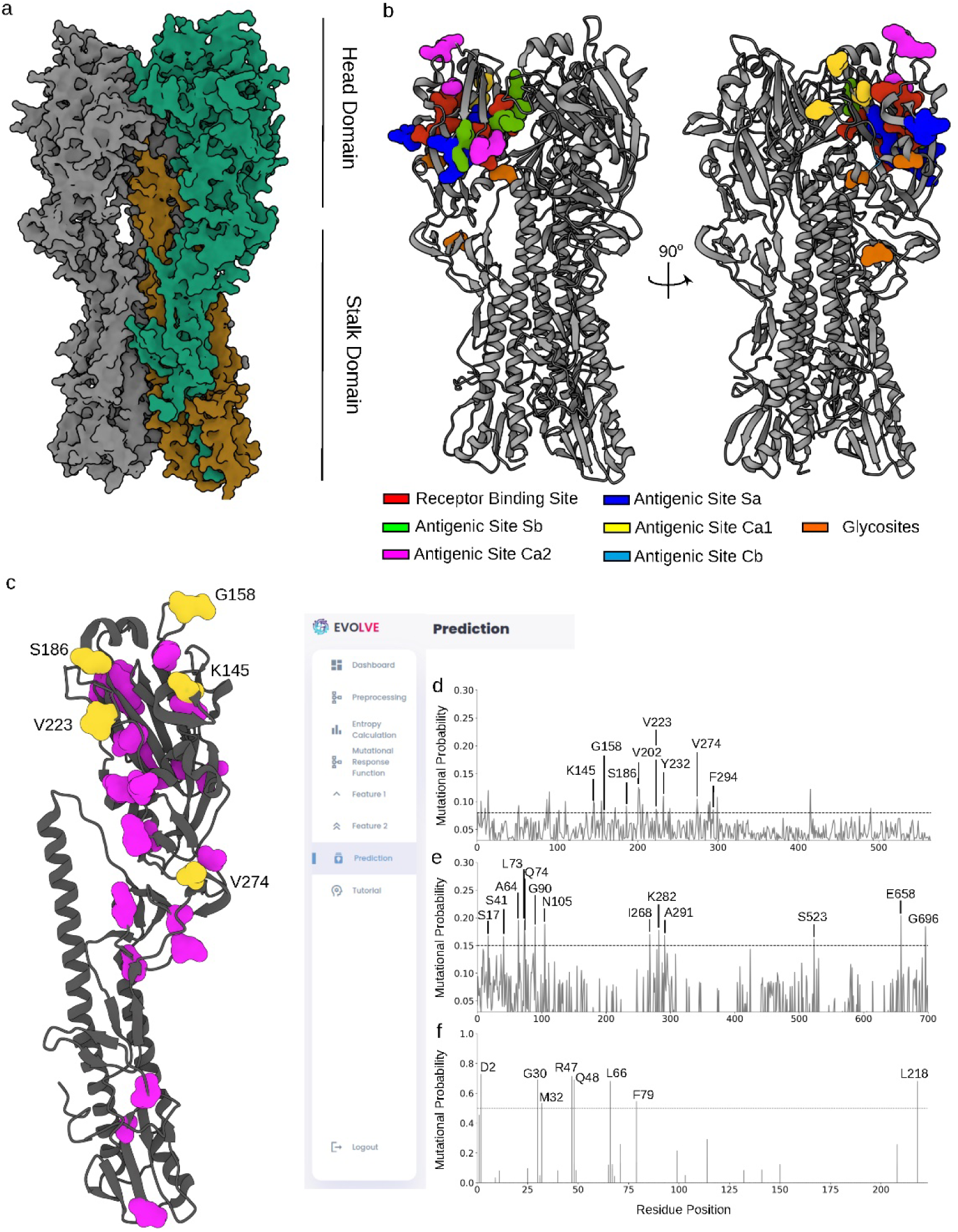
EVOLVE results for *Xanthomonas oryzae* and *Citrus Tristeza Virus*. (a) Surface view of the trimeric HA1 subunit of Hemagglutinin protein of H1N1 Influenza virus depicting head and stalk domain. (b) Cartoon representation of HA1 subunit showing import sites. (c) Single chain subunit of HA1 protein showing predicted residues (magenta) and predicted residues that have been shown to mutate (yellow with label). (d) Mutational Probability curve depicting the model prediction for influenza H1N1 HA1 subunit. (e) Mutational Probability curve depicting the model prediction for XopN protein of *Xanthomonas oryzae*. (f) Mutational Probability curve depicting the model prediction for coat protein of CTV virus.

### Case III: Xanthomonas oryzae

*Xanthomonas oryzae pv. oryzae* (Xoo) is the pathogen responsible for rice bacterial blight, a significant threat to rice cultivation^57^. Among the major virulence factors secreted by Xoo is XopN, a Type III Secretion System (T3SS) effector protein. XopN plays a critical role in manipulating the defense mechanisms of rice plants, making it essential for the pathogenicity of Xoo. Studies have shown that the deletion of XopN significantly reduces the bacterium’s ability to infect rice cultivars expressing OsSWEET11, a sugar transport protein, suggesting that XopN enables Xoo to circumvent host defenses^58^. Moreover, XopN interacts with specific rice proteins, such as OsVOZ2 and a putative thiamine synthase, enhancing its virulence^59^. Given its importance in pathogenicity and its potential for significant economic impact on rice production, XopN is a compelling target for mutation prediction. Our analysis identified 41 potential mutation sites (Supplementary Table S3) with a predicted mutational probability above the threshold of 0.12 (Fig. 5e, Table 2). We also evaluated the stability of various mutations in XopN, discovering several mutation combinations that may confer increased stability to the bacterial protein (Supplementary Fig. S5). This analysis streamlines the identification of key mutants for future experimental studies by significantly reducing the search space for relevant residues.

**Table 2:** Model prediction for potential mutating residues in CTV and *Xanthomonas oryzae* above threshold.

| CTV Mutation Prediction |  | *X. oryzae* Mutation Prediction |  |
| --- | --- | --- | --- |
| Position | Residue | Position | Residue |
| 2 | Aspartic Acid | 31 | Histidine |
| 30 | Glycine | 88 | Asparagine |
| 32 | Methionine | 91 | Asparagine |
| 47 | Arginine | 106 | Leucine |
| 48 | Glutamic Acid | 524 | Aspartic Acid |
| 66 | Lysine | 531 | Isoleucine |
| 79 | Phenylalanine | 581 | Proline |
| 218 | Leucine |  |  |

### Case IV: Citrus Tristeza Virus (CTV)

To investigate the genetic diversity among *Citrus Tristeza Virus* (CTV) isolates, several genomic regions have been identified and analyzed^60,61,62^. For this study, we focused on the Coat Protein (CP) of the virus, a multifunctional protein critical for viral structure, replication, movement, and host interaction^62^. EVOLVE predicted eight potential mutation sites exceeding the threshold cutoff of 0.5, indicating a higher likelihood of future mutations (Fig. 5f). Further analysis confirmed that mutations have already occurred at four of these sites (positions 2, 30, 32, and 48). Detailed positional information is provided in Table 2.

#### Leveraging Transfer Learning: Enhancing Model Performance through Knowledge Transfer

The extensive training required for EVOLVE and the frequent scarcity of comprehensive sequential data for many pathogens present a significant challenge. To address this, we introduce a transfer learning approach (Fig. 6a) that leverages our pre-trained model, allowing for rapid adaptation to new pathogens with limited available data. We showcase this methodology (see Method section) using the Dengue virus NS3 protease and glycoprotein of Ebola virus as a case study. By fine-tuning our SARS-CoV-2-trained model on Dengue dataset, and Influenza-trained model on Ebola dataset, we achieved remarkable predictive accuracy for potential mutation sites in the NS3 protease. This transfer learning strategy not only dramatically reduces the computational resources and time required for training but also extends EVOLVE’s utility to a broader range of pathogens, potentially accelerating our response to emerging infectious threats.

**Fig. 6:**
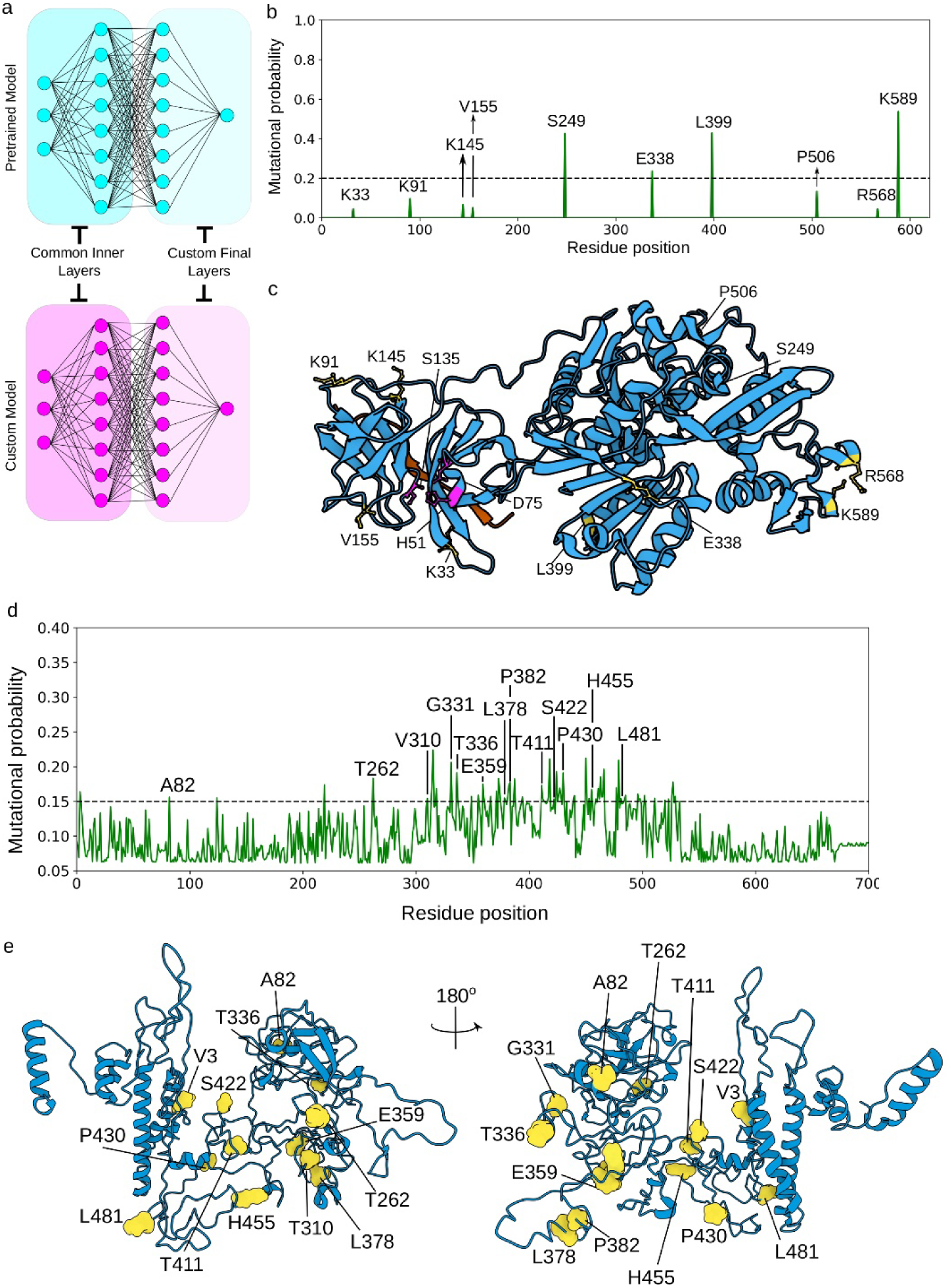
Transfer Learning Approach. (a) Schematic representation of transfer learning showing the common and custom layers of the pre-trained model and custom model. (b) Mutational Probability curve for NS3 protein of Dengue virus. (c) Mapping predicted residues on the NS3 protein (PDB ID: 2VBC) of Dengue virus depicting NS3 domain (cyan) and partial polyprotein for NS2A and NS2B (orange). The predicted residues are highlighted in yellow (d) Mutational probability curve for glycoprotein of Ebola virus. (e) Mapping the predicted residues (yellow) on the glycoprotein (PDB ID: 3CSY) of Ebola virus depicting envelope glycoprotein GP1 (cyan)

### Case V: Dengue Virus - NS3 Protease

The transfer learning approach applied to the Dengue virus (DENV) NS3 protein identified ten potential mutational hotspots: Lys33, Lys91, Lys145, Val155, Ser249, Glu338, Leu399, Pro506, Arg568, and Lys589 (Fig. 6b). These predictions, derived from a model pre-trained on the Omicron variant of SARS-CoV-2, highlight residues of significant biological relevance within the NS3 protein structure.

#### Lys91

Lys91 is implicated in binding Panduratin A, a DENV NS2B-NS3 protease inhibitor^63^. The interaction between Lys91 and Panduratin A via ionic bonds suggests that mutations at this site could hinder antiviral efficacy.

#### Lys145 and Val155

Lys145 maintains protease structural stability by forming a salt bridge with Glu93 and hydrogen bonds with substrate residues^64^, while Val155 contributes to substrate recognition through hydrophobic interactions^65^, both critical for the protease’s activity and potential inhibitor design^66^.

Although Ser249, Glu338, Leu399, Pro506, Arg568, and Lys589 (Fig. 6c) are less characterized, their identification as mutational hotspots suggest they may play previously underappreciated roles in protease dynamics. Mutation stability analysis revealed that Lys145 mutating into Glutamic Acid (Glu) particularly stabilizes the protease, while mutations at Lys91, and Val155 (Fig. 6c, Fig. S6) can either stabilize or destabilize the enzyme, underscoring their dynamic roles. Understanding these mutations is essential for antiviral strategies, as stabilizing mutations could be leveraged for inhibitor design, while destabilizing mutations could impair viral viability.

### Case VI: Ebola Virus – Glycoprotein

EVOLVE identified 68 potential mutational hotspots in the Ebola virus glycoprotein, demonstrating remarkable alignment with experimentally observed mutations from comparative analyses between historical and contemporary epidemics. The model successfully predicted several positions (262, 310, 331, 336, 359, 378, 411, 422, 430, 455, and 481, Fig. 6d) that were subsequently found to be fixed mutations (>90% frequency) in recent Ebola outbreaks^67^. Notably, the model identified position 82, corresponding to the crucial A82V mutation (Fig. 6e) in the GP1 receptor-binding region that emerged and became fixed during recent epidemics. Similarly, the predicted position 382 aligns with the observed P382T mutation in the GP1 mucin-like region^67^. The model also highlighted several positions (378-383 cluster) proximal to this functionally significant region, suggesting potential evolutionary pressure in this domain. The predictions encompass multiple functionally critical regions, including positions near the neutralizing antibody-binding site (positions 477-485 cluster). This cluster’s proximity to position 503, where an Ala-to-Val mutation causes significant backbone distortion affecting antibody recognition^67^, underscores the model’s capability to identify evolutionarily significant regions. These findings demonstrate the model’s robust predictive power in identifying both experimentally validated mutations and novel positions of potential biological significance. The alignment between predictions and empirically observed mutations validates the model’s utility for anticipating evolutionary trajectories in viral proteins, with implications for therapeutic development and outbreak preparedness. Moreover, stability analysis revealed that substitutions at position T262 resulted in significant enhancement of the glycoprotein’s thermodynamic stability (Fig. S7). These findings suggest that mutations at this position may confer structural advantages that could influence viral fitness and persistence.

#### Limitations to be aware of

Despite its strengths, the EVOLVE-based approach does have limitations that warrant careful consideration.

1. One significant and usual limitation is its reliance on large datasets for training, which may not be available for all protein sequences, potentially limiting its applicability to less well-studied proteins. Predictions may not be as robust for less-studied diseases with sparse sequence data because the model might overfit the data, capturing noise instead of significant patterns.
2. In viral evolution, epistatic interactions are frequently observed. Currently, the model handles mutations separately, which may cause it to overlook important interactions between residues that work together to affect viral behaviour and fitness.
3. The model may not generalise well to infections with drastically different evolutionary pressures or mutation rates, even though EVOLVE has shown application to a number of viruses. For example, high-mutation-rate RNA viruses, like SARS-CoV-2, are very different from DNA viruses, and further model tuning may be necessary to get reliable predictions.
4. While MRF can quantify the fluctuation of mutational entropy at each residue as a sequence evolves, the absence of a continuous evolutionary path from parent to daughter sequences, or the presence of branched phylogenies, can obscure the mean value and mean square fluctuations of mutational entropy within each domain. However, when continuous lineage tracking is possible, these fluctuations can reveal significant functional changes, even if mutations occur at only a few sites.
5. Additionally, the ML model’s predictions, while robust, are not infallible and should be interpreted as probabilities rather than certainties.
6. Currently, EVOLVE does not predict mutations for RNA system. However, this constraint presents an opportunity for future adaptation and enhancement of the model.

The complexity of biological systems means that even the most sophisticated models can occasionally miss critical residues or overestimate the importance of certain residues. However, these limitations do not diminish the overall value of the model. Instead, they highlight the need for complementary experimental validation. The public availability of such a model is crucial, as it democratizes access to advanced analytical tools, enabling researchers worldwide to leverage computational predictions to guide their experimental designs. By integrating our model into their workflows, experimentalists can more efficiently pinpoint key mutations for further study, thereby accelerating the pace of discovery and enhancing our understanding of protein function and stability.

## Conclusion

To address the rapid evolution of viruses and pathogens, innovative tools like EVOLVE are indispensable. Leveraging the concepts of statistical mechanics of dynamic phase transition, EVOLVE analyzes the collective behavior of mutations across extensive genomic sequence spaces, while machine learning processes this vast data to extract insights at a finer scale, such as predicting site-specific mutational hotspots or identifying mutation prone domain that could significantly alter evolutionary dynamics. Together, these approaches provide powerful tools for advancing our understanding of viral adaptation and dissemination, playing a critical role in mitigating the risks posed by emerging strains.

Given the vast sequence space and the sparse occurrence of mutations, EVOLVE serves as a first stage filter, narrowing the search to a manageable set of high-priority mutations. These prioritized mutations can then undergo detailed analysis through experimental assays or computational tools to assess their effects on protein stability, functionality, and other properties. The seamless integration of EVOLVE with downstream methodologies enhances pipeline efficiency, enabling focused validation and a proactive approach to studying and anticipating viral evolution.

This communication shows that EVOLVE’s application extends beyond SARS-CoV-2 to pathogens like H1N1, Citrus Tristeza Virus, and *Xanthomonas oryzae*, demonstrating its versatility in pathogen surveillance. Through a transfer learning approach, EVOLVE can also predict mutations in understudied pathogens with limited data, offering insights for emerging diseases.

Looking ahead, EVOLVE’s modular architecture enables continuous improvement and expansion. Planned updates include integrating predictions with AlphaMissense^68^—a tool based on AlphaFold^69^ that assesses missense variant pathogenicity using population frequency data— and incorporating advanced structure-based simulations to evaluate the molecular impact of predicted mutations, including the exploration of structural and functional dynamics in proteins from concerning variants. By uniting the sequence-structure-function paradigm, EVOLVE aims to advance drug development and vaccine design, empowering researchers to anticipate and respond to future viral outbreaks more effectively.

## Methods

### Implementation

EVOLVE streamlines sequential informatic analysis and allows user uploads, making the exploration of genomic and protein sequences easier. EVOLVE web server architecture is built on a three-tier structure: a server layer that handles data processing and communication via Flask, a client layer that renders templates using Flask, and a SQLite3 database that stores user data, authentication, and analysis results. The user interface of the web application is formed by the client layer. The core web technologies—HTML, CSS, and JavaScript—are used in its development. The Cascading Style Sheets (CSS) provide styling and layout, HTML (Hyper Text Markup Language) provides the structural framework, and JavaScript is used for client-side scripting, which allows for interactive features and form handling. The Flask web framework is essential in managing different facets of the server-side functionality at the server layer. Flask serves as the basis for using decorators to direct incoming requests to endpoints. It uses functions to control the rendering of HTML templates, making it easier for users to see dynamic content. Furthermore, Flask works in conjunction with libraries like pandas to facilitate effective data handling, and it plays a crucial role in managing file uploads and data processing. The SQLite3 database is the main data management tool and functions as the backend storage mechanism. It also serves as a repository for user-specific data, analysis results, and uploaded files, guaranteeing safe and well-organized storage.

### Server Configuration

The web application is hosted on a server environment with an economical hardware specification, including 0.15 CPU units, and 512MB of RAM. The Flask application is deployed directly on the server. By reducing the requirement for extra server layers, this strategy maximizes the hosting environment’s efficiency. The web server is configured using the Web Server Gateway Interface (WSGI) that acts as an interface between web servers and Python-based web applications.

### Model Training

Model training was optimized by evaluating various training-to-testing set ratios to identify the most effective configuration (see Supplementary Fig. S7). The analysis determined that a ratio of 85:15 provided the best model performance, balancing robust learning with accurate validation.

#### Entropy

User-supplied MSA files in the *.aln* format can be used in EVOLVE’s “***Entropy Calculation***” tab to calculate the Shannon entropy of particular sites in protein or genomic sequences. The calculation of Shannon entropy is a useful tool for assessing evolutionary variability since it clarifies how mutations cause specific sites to change over time. The entropy values of sites that show conservation over evolutionary timelines are generally lower than those of sites that show greater variability. The Shannon entropy *H_i_* of a given site *i* is computed using the formula:

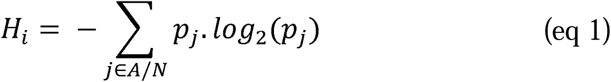

Here, *p_j_* stands for the likelihood that a particular amino acid (for proteins) or nucleotide (for genomics) will occur at that specific site, and *H_i_* is the Shannon entropy of site *i*. The range of potential amino acids or nucleotides is represented by the index *j*. *j* is a member of the protein groups *A*, where *A* = {A, C, D, E, F, G, H, I, K, L, M, N, P, Q, R, S, T, V, W, Y}, and the genomic sequence group *N*, where *N* = {A, T, G, C}. This computation provides users with useful information about the evolutionary dynamics of sequence sites by emphasizing the conservation or variability linked to mutations over time. Users can calculate the entropy for parent and offspring variants to analyze the evolution of the sequence of variants using mutational entropy.

#### Mutational Response Function

EVOLVE forecasts a species’ transitional trajectory in response to accumulated mutations by utilizing the predictive power of the Mutational Response Function (MRF) (see Supplementary File for more detail). MRF is a crucial metric that summarizes the chronology of species transitions and clarifies the evolutionary trajectory of the species. The monthly organized entropy data is presented in a CSV format, which is the input data needed for the MRF computation. It is convenient for users to upload this formatted data file onto the EVOLVE interface. Following examination, EVOLVE processes the data, determines the MRF, and produces an output that displays the MRF values. Users can supply the mutational entropy data of ancestor and offspring variants to capture the transition point in the evolution timeline.

#### Mutation Prediction

EVOLVE allows the user to predict the potential mutational sites in a protein sequence of the species. The user can calculate the features for model training by visiting the “*Feature 1*” and “*Feature 2*” (see Supplementary File section “Feature selection for Model Training”) tab in the dashboard. “*Feature 3*” of the model is the entropy of the specific sites that can be calculated using the “***Entropy Calculation***” tab. A detailed explanation of individual features and how to prepare the data is provided in the form of a tutorial at https://evolve-0fm3.onrender.com/theory_corner. After compiling the dataset, the user can upload the data file in the “*Prediction*” tab. This will allow the user to utilize the Multi-Layer Perceptron (MLP) model for the prediction of potential mutating residues in the protein sequence. These calculated values provide users with informative information, providing the basis for graphing and visual aids, allowing a thorough representation of the evolutionary history of the species over time.

### Transfer Learning

The EVOLVE web server provides a robust framework for users to apply pre-trained models to new viral sequences through a sequence similarity check, designed to enhance the accuracy and relevance of mutation predictions. This feature addresses two critical scenarios for researchers working with novel sequences. First, if a user wishes to leverage any of the pre-trained models provided by EVOLVE, they can initiate a sequence similarity search against the pre-existing models hosted on the platform. Currently, EVOLVE offers four pre-trained models specifically tailored to the Omicron variant of SARS-CoV-2, XopN of *Xanthomonas oryzae*, Coat Protein Citrus Tristeza Virus (CTV), and the Hemagglutinin protein H1N1 Influenza virus. The web server will compute a cosine similarity score between the user-provided sequence and these available models. The cosine similarity for two protein sequences is defined as:

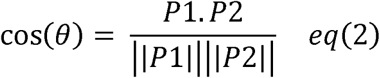

Where P1 and P2 correspond to two protein sequences represented as vectors:

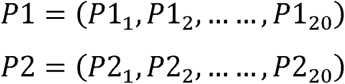

Where *P*1*_i_* and *P*2*_i_* represent the frequency of the *i^th^* amino acid in proteins P1 and P2 respectively. Equation 2 can be expanded for protein sequences such as:

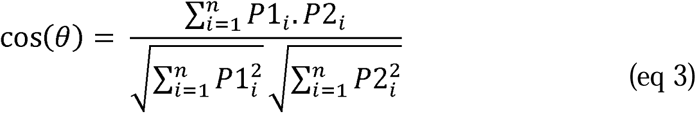

Where, Σ□ denotes the sum from i=1 to 20 for the 20 standard amino acids.

If the similarity score exceeds 0.85, the system recommends using the corresponding pre-trained model for mutation prediction. However, it is advised that any predictions generated should be further validated through experimental methods to ensure accuracy. For cases where the sequence similarity scores are below the 0.85 threshold across all available models, or if the user prefers not to use a pre-trained model, EVOLVE offers a transfer learning approach. Users are encouraged to access our GitHub repository, where pre-trained versions of the model are available for download. These models can then be utilized in transfer learning, allowing users to fine-tune the model on their specific dataset. In this approach, the initial layers of the model remain frozen to preserve the learned features from the original training, while the final layers are adjusted to accommodate the new sequence data, thus creating a tailored model for accurate mutation prediction.

Prior to applying transfer learning method for Dengue virus and Ebola virus, a similarity score checks between the protein sequences of SARS-CoV-2 and Dengue virus, and SARS-CoV-2 and Ebola virus was conducted to ensure the applicability of the transfer learning approach. Only protein sequences exhibiting a similarity score above 0.85 were selected for further analysis, ensuring that the model’s learned features would be relevant and transferable. In the transfer learning process, we strategically froze the initial layers of the model, which capture fundamental features from the Omicron dataset, while allowing the final layers to remain unfixed. This design enabled the model to adapt to the unique characteristics of the Dengue virus dataset, while retaining the generalized knowledge acquired from the Omicron dataset. By keeping the foundational layers intact and fine-tuning the higher layers, the model could effectively integrate new data without losing the crucial patterns identified during its initial training.

## Data Availability

The requisite codes and data files are available at https://www.github.com/psychedelic2007/Evolve. The repository also provides example files for the user to understand how the data file must be prepared and supplied.

## Code Availability

The Python codes for local calculation are freely available at https://www.github.com/psychedelic2007/Evolve. EVOLVE is freely accessible at https://www.evolve-0fm3.onrender.com.

## Author Contributions

Conceptualization: SR; Data curation, Modelling, and Server Development: SS; Formal analysis: SS, SR, AS, MBN; Funding acquisition: SR; Methodology: SR, SS, RS; Project administration and Resources: SR; Validation: SS, SR; Writing – original draft: SS, SR; Writing – review & editing: SR, SS, AS, AM, MBN

## Acknowledgements

SR acknowledges support from the Department of Biotechnology (DBT) (Grant No. BT/12/IYBA/2019/12 and BT/PR40192/BTIS/137/692023).

