## Supplementary file for "EVOLVE: A Web Platform for Evolutionary Phase Analysis and New Variant Exploration from Multi-Sequence Data"

**This Supplementary File includes:**

- Lasso Regression
- Fig S1 showing the feature importance, using Lasso regression, between old and updated model
- Table S1 shows the variant analysis for the four major predicted mutating positions in the Omicron variant
- Fig S2 showing Omicron Mutational Analysis
- Fig S3 showing residue distribution in Omicron with only entropic consideration versus ML-assisted predictions
- Fig S4 showing Influenza Mutational Analysis
- Table S2 shows the variant analysis for the predicted mutating position in *Xanthomonas oryzae*
- Fig S5 showing *Xanthomonas* Mutational Analysis
- Fig S6 showing Dengue Mutational Analysis
- Fig S7 showing Ebola Mutational Analysis
- Table S4 shows the variant analysis for the predicted mutating position in glycoprotein of Ebola virus
- Fig S8 shows the ROC-AUC curve for the different trains to test ratio
- Site Specific Entropy
- Mutational Response Function
- Feature Selection for Model Training
- Importance of ML model over pure Entropic Calculation

**Lasso Regression**

One technique used to deal with the problem of model overfitting is regularization. It entails the cost function's insertion of coefficient terms, or betas, which penalize and limit these terms to lesser magnitudes. This method helps to capture important trends in the data while also avoiding overfitting, which happens when the model gets too complicated. Regularisation basically aims to avoid the extremes of excessive complexity and excessive simplicity by maintaining the model's simplicity while ensuring that it performs well on the training set.

In the context of Linear Regression, with a dataset comprising N samples $\left\{ \left( x_{i},y_{i} \right) \right\}N_{i}=1$, where each $x_{i}=(x_{i1},\ldots.,x_{ip})$ represents a p-dimensional vector of features, and each $y_{i}\in R$ denotes the associated response variable. The primary aim is to estimate the response variable $y_{i}$ by employing a linear combination of these features.

*(eq S1)*

$$\eta\left( x_{i} \right)= \beta_{0}+ \sum_{j=1}^{p} x_{ij}\beta_{j}$$

For determining the best fit, the associated cost function requires optimization. In the case of Linear Regression, the prevalent cost function employed is the Mean Squared Error (MSE):

*(eq S2)*

$$\arg\min\left\{ \frac{1}{N}\sum_{i=1}^{N} \left( y_{i}- \beta_{0}- \sum_{j=1}^{p} x_{ij}\beta_{j} \right)^{2} \right\}$$

In Lasso regression, an additional regularization term, involving the "*sum of the absolute values of the coefficients*," is incorporated into the previously mentioned cost function. The Lasso method effectively shrinks the coefficients of irrelevant variables to zero, consequently facilitating direct feature selection as part of the modelling process.

*(eq S3)*

$$\arg\min\left\{ \frac{1}{N}\sum_{i=1}^{N} \left( y_{i}- \beta_{0}- \sum_{j=1}^{p} x_{ij}\beta_{j} \right)^{2} \right\}+ \lambda\sum_{j=1}^{p} |\beta_{j}|$$

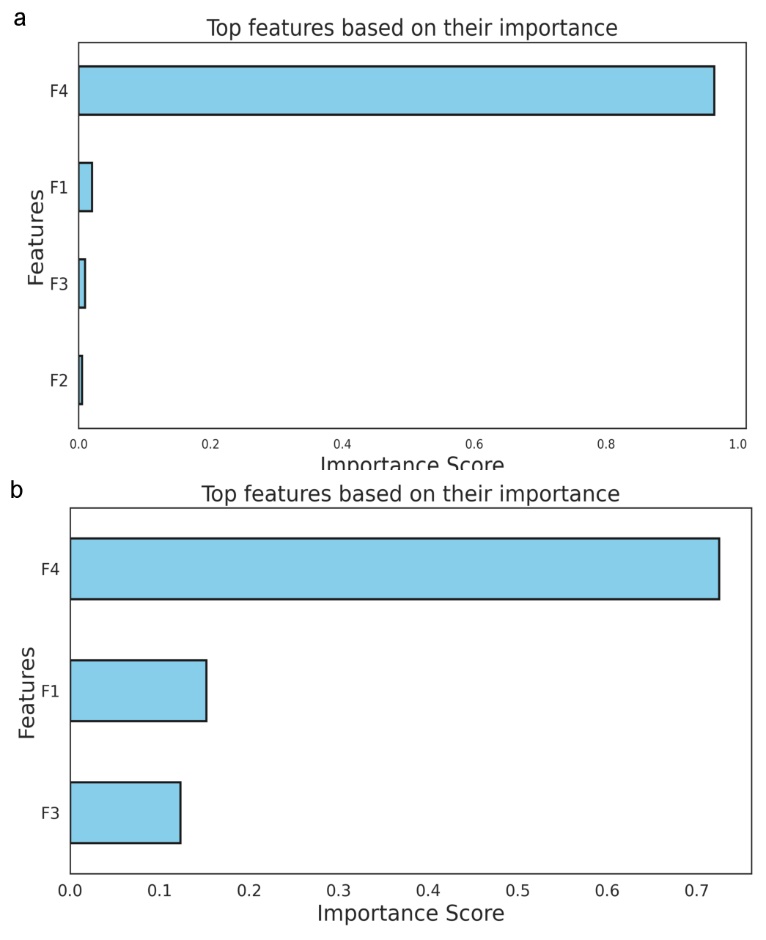

**Fig. S1: Feature importance analysis using LASSO regression.** (a) When the model used four features^1^, the fourth feature i.e. “Entropy” contributed more towards model learning. Feature 2 i.e. “Distribution Probability” didn’t contribute much towards model training. (b) With three features the feature importance of Feature 1 and Feature 3 also increases but Feature 4 remains the dominant feature via which the model learns.

**Table S1:** Variant analysis for the four major predicted mutating position in Omicron variant

| Variant Name | N334 | S494 | A653 | T791 |
| --- | --- | --- | --- | --- |
| BJ.1 | N334K (USA), N334S (Spain), N334Y (USA) | S494P (Kenya, Slovakia) | A653V (Portugal, USA) | T791A (India, Portugal, USA) |
| XBB.1.5.68.mix | - | S494A (UK) | - | - |
| XBB.1.16.1 | - | S494A (USA) | A653V (USA) | - |
| EG.5.1.1 | - | S494P (Japan) | A653V (Japan) | - |
| FY.4.1.2 | - | S494P (Kenya) | - | - |
| XBB | - | S494P (Kenya) | A653V (UK) | - |
| FY.4.1 | - | S494P (Kenya) | - | - |
| XBB.1.5 | - | - | A653T (UK, USA) | T791I (UK) |
| BQ.1.1.29 | - | - | A653V (Portugal) | - |
| BA.4.6.5 | - | - | A653V (Switzerland) | - |
| BA.2.38 | - | - | A653V (UK) | - |
| B.1.1.529 | - | - | A653V (UK) | T791I (USA) |
| DU.1 | - | - | A653V (UK) | - |
| XBB.1.5.4 | - | - | A653V (USA) | T791I (USA) |
| BA.2.12.1 | - | - | A653V (USA) | - |
| EG.5.1.6 | - | - | A653V (USA) | - |
| BA.2 | - | - | A653V (USA) | - |
| EG.5.1 | - | - | A653V (USA) | - |
| HZ.3 | - | - | A653V (USA) | - |
| BA.2.3.20 | - | - | A653V (USA) | - |
| BF.7 | - | - | A653V (USA) | - |
| BA.5.2.1 | - | - | A653V (USA) | - |
| BA.5.1 | - | - | A653V (USA) | - |
| JN.2 | - | - | - | T791I (Denmark) |
| EG.1.6 | - | - | - | T791I (Japan) |
| BA.1.mix | - | - | - | T791I (Switzerland) |
| XBB.1.16.15 | - | - | - | T791I (UK) |
| BA.2.86.1 | - | - | - | T791I (UK) |
| XBB.1.9.1.mix | - | - | - | T791I (USA) |
| XBB.1.16.6 | - | - | - | T791I (USA) |
| XBB.1.42.2 | - | - | - | T791K (USA) |
| XBB.1 | - | - | - | T791K (USA) |

**Omicron Mutational Analysis**

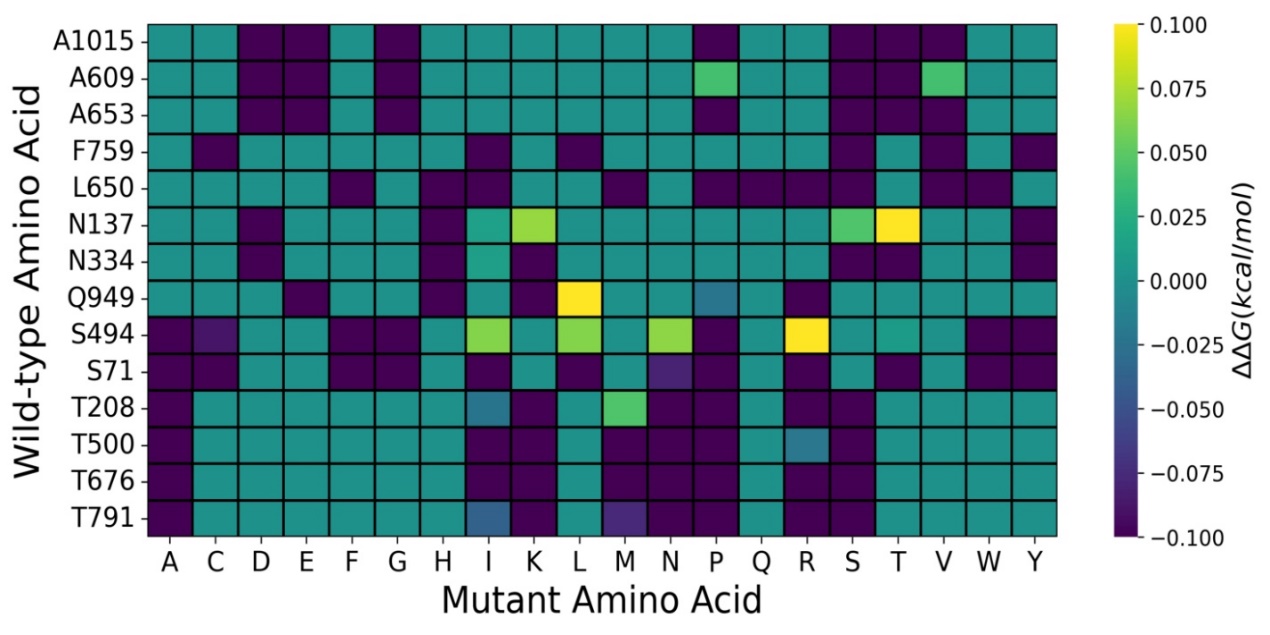

**Fig. S2: Heatmap of point mutation stability/instability analysis of Omicron.** Y axis corresponds to the predicted mutational residues by the model and x axis corresponds to the amino acids into which the predicted mutational residues can mutate into. The colorbar corresponds to the ∆∆G value. Positive value corresponds to stabilising mutation whereas negative values correspond to destabilising mutations.

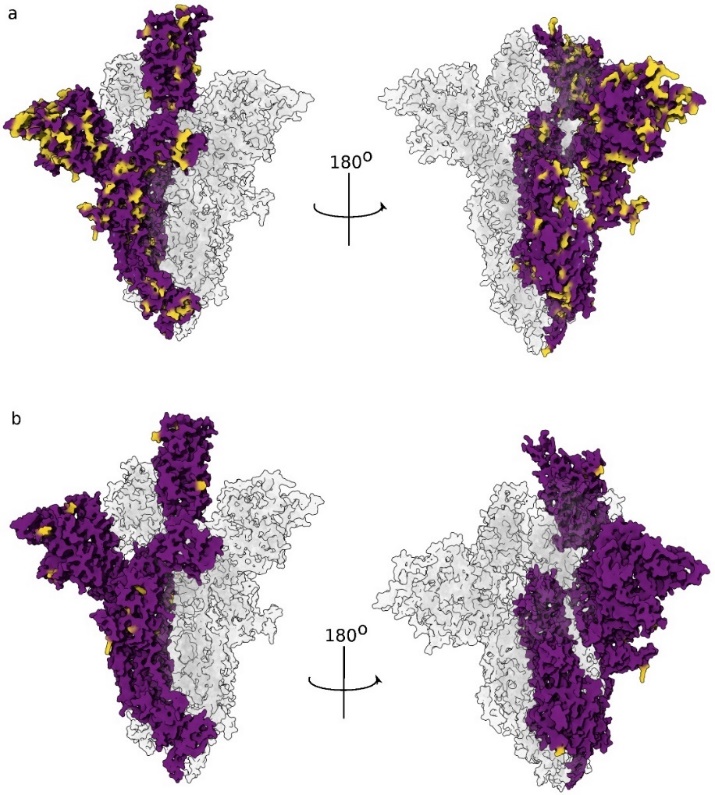

**Fig. S3: Residue distribution results for Omicron variant** (a) Residues with variable entropic values as obtained from pure entropy analysis is represented in yellow (b) Predicted residues from the ML model (yellow).

**Influenza Mutation Analysis**

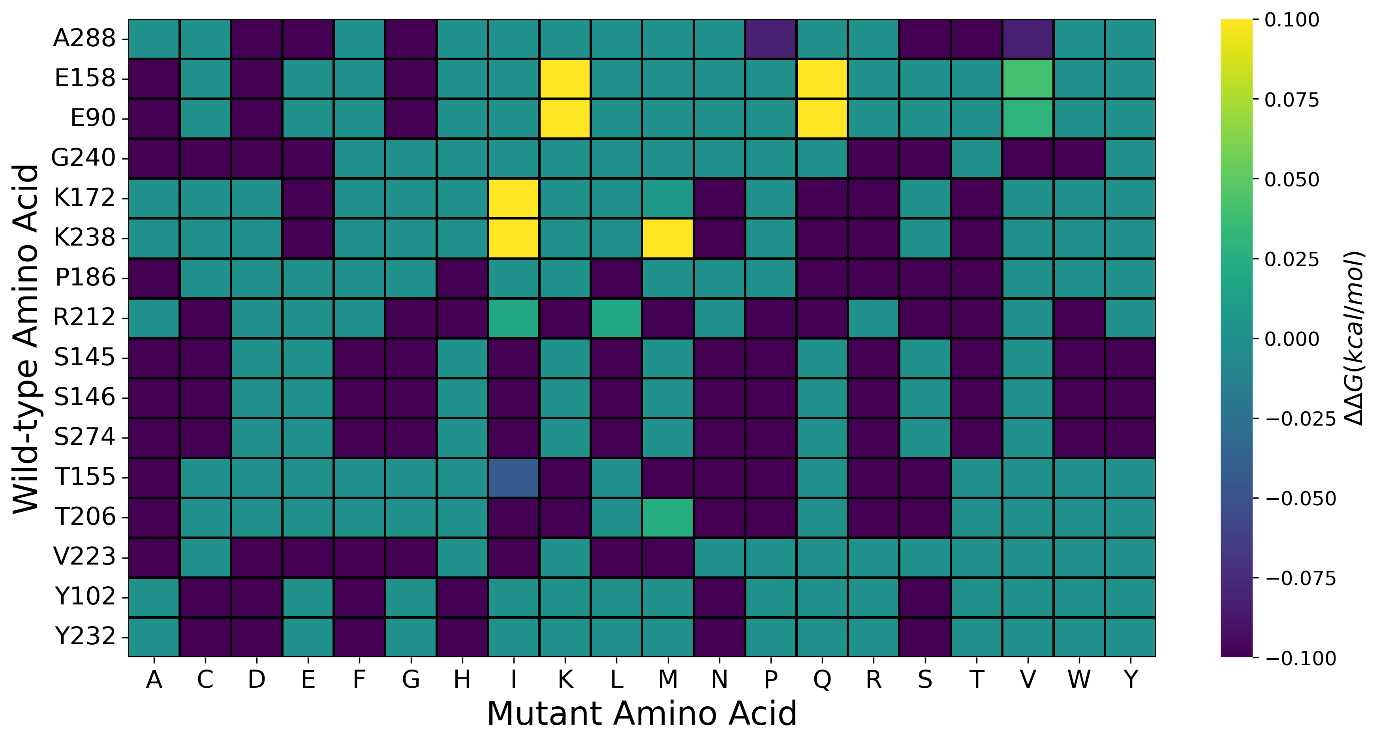

**Fig. S4: Heatmap of point mutation stability/instability analysis of Influenza H1N1 protein.** Y axis corresponds to the predicted mutational residues by the model and x axis corresponds to the amino acids into which the predicted mutational residues can mutate into. The colorbar corresponds to the ∆∆G value. Positive value corresponds to stabilising mutation whereas negative values correspond to destabilising mutations.

**Table S2:** Variant analysis for the predicted mutating position in HA1 Influenza H1N1

| Position | Residue | Position | Residue |
| --- | --- | --- | --- |
| 14 | Cysteine | 202 | Valine |
| 52 | Cysteine | 203 | Phenylalanine |
| 87 | Isoleucine | 206 | Serine |
| 90 | Proline | 212 | Lysine |
| 102 | Phenylalanine | 223 | Valine |
| 110 | Glutamine | 232 | Tyrosine |
| 145 | Lysine | 238 | Glutamine |
| 146 | Serine | 240 | Glycine |
| 155 | Valine | 274 | Valine |
| 158 | Glycine | 288 | Isoleucine |
| 172 | Lysine | 290 | Threonine |
| 186 | Serine | 294 | Phenylalanine |
| 201 | Tyrosine | 299 | Proline |

**Table S3:** Variant analysis for the predicted mutating position in *Xanthomonas oryzae*

| Position | Residue | Position | Residue |
| --- | --- | --- | --- |
| 8 | Threonine | 121 | Threonine |
| 10 | Leucine | 191 | Tyrosine |
| 11 | Asparagine | 212 | Phenylalanine |
| 15 | Proline | 269 | Alanine |
| 18 | Proline | 275 | Cysteine |
| 20 | Glycine | 280 | Alanine |
| 30 | Alanine | 283 | Aspartic Acid |
| 31 | Histidine | 288 | Asparagine |
| 38 | Proline | 292 | Glutamic Acid |
| 42 | Glutamine | 296 | Aspartic Acid |
| 47 | Methionine | 297 | Phenylalanine |
| 65 | Serine | 309 | Threonine |
| 67 | Alanine | 425 | Glycine |
| 74 | Glutamic Acid | 505 | Aspartic Acid |
| 75 | Serine | 524 | Aspartic Acid |
| 77 | Glutamine | 531 | Isoleucine |
| 80 | Valine | 581 | Proline |
| 87 | Proline | 659 | Alanine |
| 88 | Asparagine | 692 | Glutamic Acid |
| 91 | Asparagine | 697 | Valine |
| 106 | Leucine |  |  |

***Xanthomonas oryzae* mutation analysis**

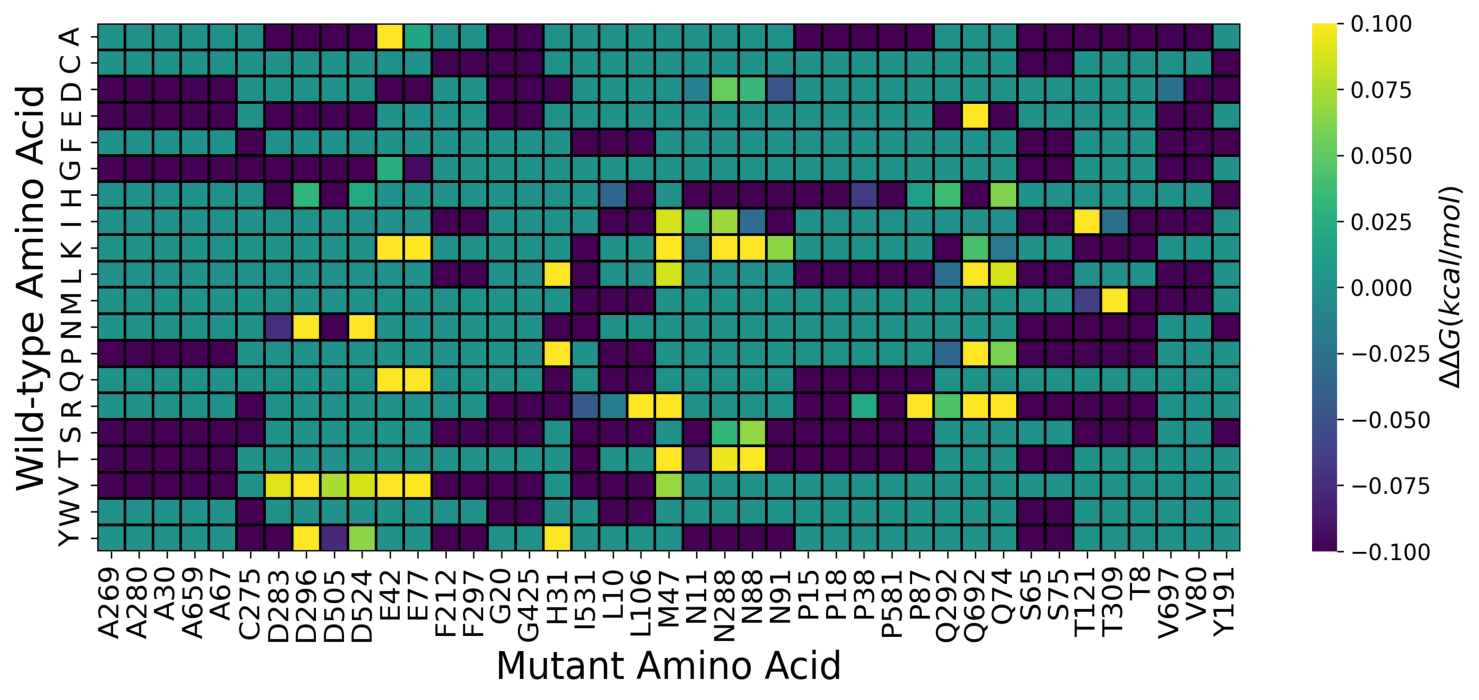

**Fig. S5: Heatmap of point mutation stability/instability analysis of XopN.** Y axis corresponds to the amino acids into which the predicted mutational residues can mutate into, and x axis corresponds to the predicted mutational residues by the model. The colorbar corresponds to the ∆∆G value. Positive value corresponds to stabilising mutation whereas negative values correspond to destabilising mutations.

**Dengue Protease Mutational Analysis**

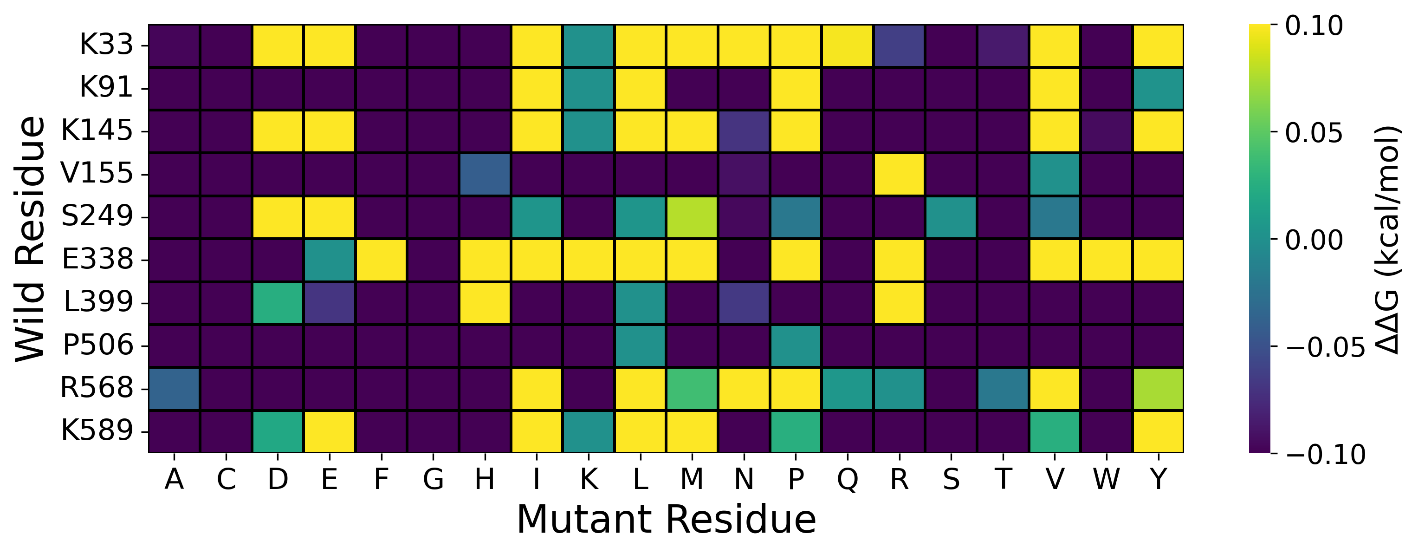

**Fig. S6: Heatmap of point mutation stability/instability analysis of NS3 protein of Dengue virus.** Y axis corresponds to the predicted mutational residues by the model and x axis corresponds to the amino acids into which the predicted mutational residues can mutate into. The colorbar corresponds to the ∆∆G value. Positive value corresponds to stabilising mutation whereas negative values correspond to destabilising mutations.

**Ebola Glycoprotein Mutation Analysis**

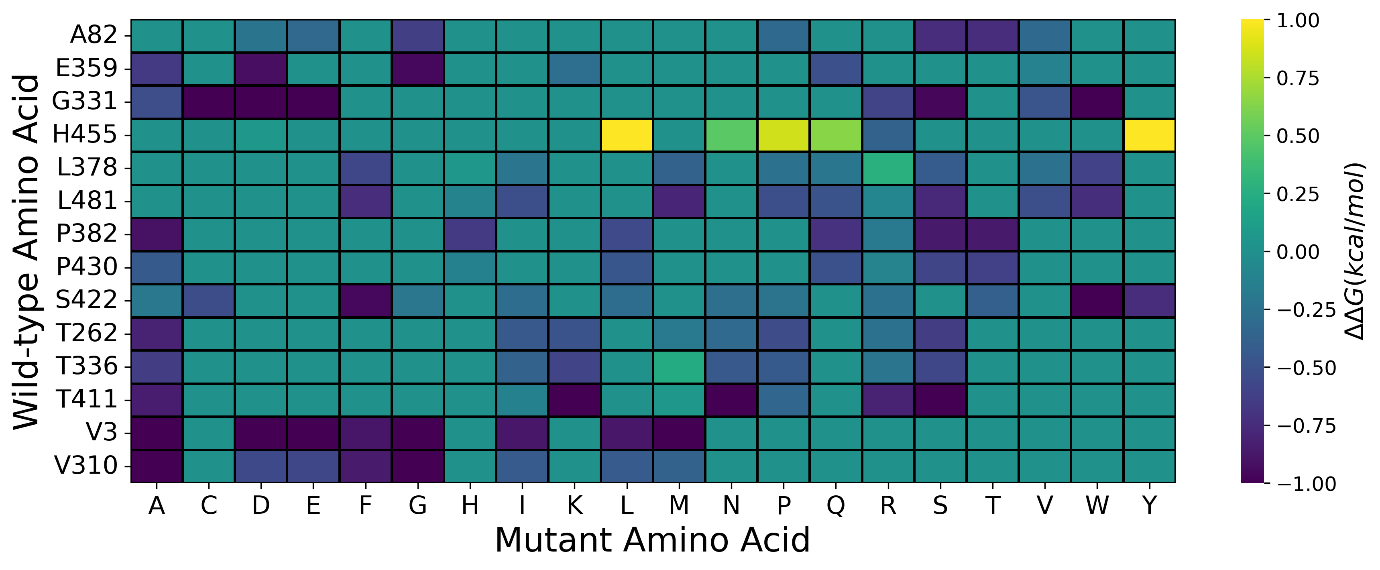

**Fig. S7: Heatmap of point mutation stability/instability analysis of glycoprotein of Ebola virus.** Y axis corresponds to the predicted mutational residues by the model (having associated literature study) and x axis corresponds to the amino acids into which the predicted mutational residues can mutate into. The colorbar corresponds to the ∆∆G value. Positive value corresponds to stabilising mutation whereas negative values correspond to destabilising mutations.

**Table S4:** Variant analysis for the predicted mutating position in glycoprotein of Ebola virus

| Position | Residue | Position | Residue |
| --- | --- | --- | --- |
| 3 | Valine | 425 | Threonine |
| 82 | Alanine | 426 | Alanine |
| 124 | Alanine | 427 | Alanine |
| 129 | Isoleucine | 428 | Glycine |
| 262 | Threonine | 430 | Proline |
| 310 | Valine | 439 | Lysine |
| 314 | Glycine | 440 | Serine |
| 315 | Alanine | 450 | Threonine |
| 318 | Isoleucine | 451 | Serine |
| 331 | Glycine | 453 | Glutamine |
| 336 | Threonine | 455 | Histidine |
| 337 | Glutamic Acid | 459 | Alanine |
| 349 | Alanine | 461 | Asparagine |
| 355 | Serine | 462 | Asparagine |
| 359 | Glutamic Acid | 463 | Asparagine |
| 360 | Alanine | 464 | Threonine |
| 366 | Threonine | 465 | Histidine |
| 371 | Isoleucine | 466 | Histidine |
| 373 | Threonine | 477 | Glycine |
| 374 | Serine | 479 | Leucine |
| 378 | Leucine | 481 | Leucine |
| 379 | Threonine | 484 | Asparagine |
| 381 | Lysine | 485 | Threonine |
| 382 | Proline | 496 | Glycine |
| 383 | Glycine | 497 | Arginine |
| 387 | Serine | 498 | Arginine |
| 394 | Tyrosine | 502 | Glutamic Acid |
| 411 | Threonine | 511 | Cysteine |
| 412 | Aspartic Acid | 520 | Threonine |
| 413 | Asparagine | 523 | Glutamic Acid |
| 417 | Alanine | 526 | Alanine |
| 418 | Serine | 527 | Isoleucine |
| 422 | Serine | 529 | Leucine |
| 424 | Threonine | 533 | Proline |

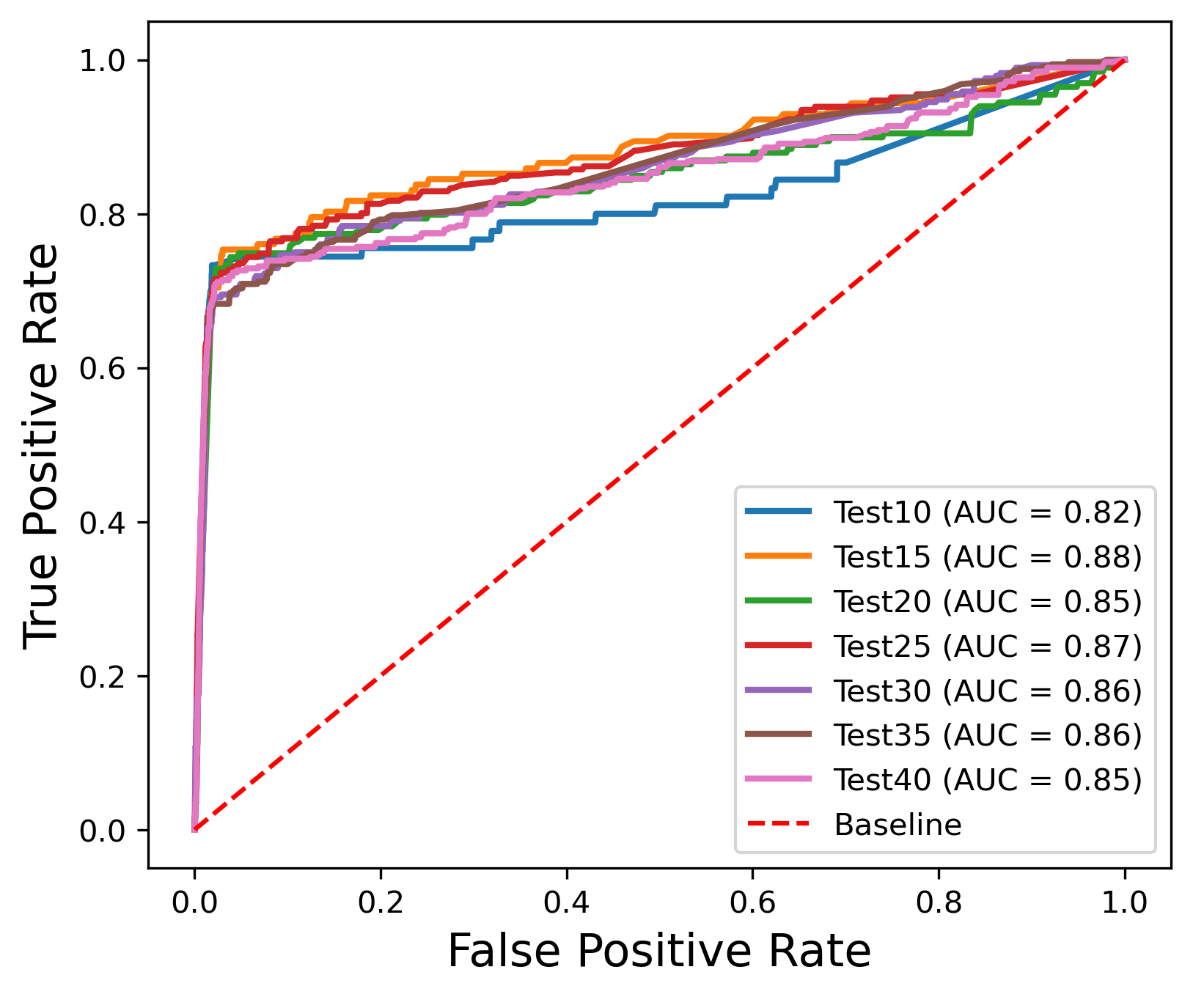

**Fig. S8: ROC analysis of the ML model.** The analysis revealed that having a balance of 85% training and 15% test files (85:15, yellow curve AUC=0.88) showed the best area under the curve thus recommending the usage of a similar ratio of train: test files.

***Mutational Response Function***

A strong mathematical foundation that takes into consideration the dynamic changes that happen at each residue along the sequence is necessary for the construction of the Mutational Response Function (MRF). This paradigm is based on the idea of entropy fluctuation, in which the entropy of every residue is monitored and measured over the course of the virus's evolution. The mutational response of each residue is carefully measured by the MRF, which captures the intrinsic diversity and adaptability of the viral genome by following the sequence evolution over a predetermined temporal interval. Mathematically MRF can be represented as:

$$MRF\left( i \right)= \frac{\left\langle\Delta S{(i)}^{2} \right\rangle}{\bar{S}(i)}= \frac{\frac{1}{T}\sum_{t=1}^{T} {(S\left( i \right)- \bar{S})}^{2}}{\bar{S}} eq(S4)$$

Where, $S(i)$ is the entropy of the $i^{th}$ nucleotide/amino acid, $\bar{S}$ is the time-averaged entropy of that $i^{th}$ nucleotide/amino acid for 3 months (T = 3).

**Feature Selection for Model Training**

***Feature 1: Pair Predictability of Amino Acids***

This feature relies on the fact that different amino acids occur in different pairs in a protein sequence and a mutation in one amino acid can possibly change the amino acid pair resulting in a different pair frequency. Based on the occurrence of these variable pairs for a particular amino acid, we can calculate the actual frequency and the predicted frequency of the pairs as:

$$\frac{No. of "x"}{L}X\frac{No. of "y"}{L-1}X\left( L-1 \right) eq(S5)$$

Where,

$x$ is the first amino acid of interest and $y$ is the second amino acid of interest making the pair $xy$, and L is the length of the sequence.

***Feature 2: Distribution Probability of Amino Acids***

This feature is based upon the occupancy of sub-populations and partitions. In a protein sequence, certain amino acids cluster together in specific regions rather than homogeneously spreading throughout the length of the protein. Thus, any change/mutation in a particular amino acid will result in the change of how that amino acid is distributed in a protein sequence. The position of each type of amino acid in a protein sequence can be viewed as a particular distribution whose probability can be calculated as:

$$Distribution Probability= \frac{r!}{(q_{o}!X q_{1}!X\ldots Xq_{n}!)}X\frac{r!}{(r_{1}!Xr_{2}!X\ldots Xr_{n}!)}Xn^{-r} eq(S6)$$

In this case, $n$ denotes the number of partitions, $r_{n}$ is the number of amino acids in the $n^{th}$ partition, and $q_{n}$ denotes the number of partitions that have the same number of amino acids. The fundamental particles in statistical mechanics are arranged in the energy state with reference to this distribution probability.

***Feature 3: Future Count of Amino Acid***

This feature is dependent on the translation probabilities between the translated amino acids and the RNA codons. This characteristic includes the possibility of an amino acid mutating into another amino acid, or the mutational probability of an amino acid. The fact that codon codes for an amino acid serves as evidence for this characteristic. A distinct codon that translates into a different amino acid is produced by any mutation in one of the nucleotide bases. This can be calculated as:

$$Future Composition= \frac{Predicted Contribution}{Actual Contribution} eq(S7)$$

***Feature 4: Amino Acid Entropy***

This feature makes it possible to quantify the impact of mutation in terms of the entropy of certain amino acid residues. The selection of this characteristic was justified by the evolutionary conservation of residue position. The residue's entropy will be zero if mutations in the protein sequence do not occur throughout numerous generations, preserving the amino acid at that specific location. However, any insertion of a mutation would alter the pattern of amino acids, raising the entropy value.

**Importance of ML model over pure Entropic Calculation**

According to the LASSO regression analysis, entropy emerges as the most significant factor during model training, holding the highest weightage among all model features. This prominence of entropy might prompt a question: why not use entropy alone to interpret the results? Given the substantial influence of entropy in identifying mutational hotspots, it might seem sufficient to base our analysis solely on entropic values. However, despite the clear importance of entropy, there are compelling reasons to integrate machine learning (ML) approaches into our methodology. When analysing sequences for mutational hotspots using entropic factors, we encounter a substantial number of residues exhibiting significant entropy (Fig. S3a). This abundance complicates the selection process of the most crucial sites for further analysis. Entropy, which measures the variability or uncertainty at specific sites in a sequence, indicates regions prone to mutations. However, the sheer volume of high-entropy residues can make it challenging to streamline and prioritize these sites effectively.
